# Forgetful inference in a sophisticated world model

**DOI:** 10.1101/419317

**Authors:** Sanjeevan Ahilan, Rebecca B. Solomon, Yannick-André Breton, Kent Conover, Ritwik K. Niyogi, Peter Shizgal, Peter Dayan

**Affiliations:** Gatsby Computational Neuroscience Unit, University College London, London, United Kingdom; Center for Studies in Behavioral Neurobiology, Concordia University, Montreal, Canada; Department of Experimental Psychology, University of Oxford, Oxford, United Kingdom

## Abstract

Humans and other animals are able to discover underlying statistical structure in their environments and exploit it to achieve efficient and effective performance. However, such structure is often difficult to learn and use because it is obscure, involving long-range temporal dependencies. Here, we analysed behavioural data from an extended experiment with rats, showing that the subjects learned the underlying statistical structure, albeit suffering at times from immediate inferential imperfections as to their current state within it. We accounted for their behaviour using a Hidden Markov Model, in which recent observations are integrated with the recollections of an imperfect memory. We found that over the course of training, subjects came to track their progress through the task more accurately, a change that our model largely attributed to decreased forgetting. This ‘learning to remember’ decreased reliance on recent observations, which may be misleading, in favour of a longer-term memory.

**Author summary:** Humans and other animals possess the remarkable ability to find and exploit patterns and structures in their experience of a complex and varied world. However, such structures are often temporally extended and latent or hidden, being only partially correlated with immediate observations of the world. This makes it essential to integrate current and historical information, and creates a challenging statistical and computational problem.

Here, we examine the behaviour of rats facing a version of this challenge posed by a brain-stimulation reward task. We find that subjects learned the general structure of the task, but struggled when immediate observations were misleading. We captured this behaviour with a model in which subjects integrated evidence from their observations together with a memory whose imperfections accounted for their errors.

The subjects’ performance improved markedly over successive sessions, allowing them to overcome misleading observations. According to the model, this arose from a process of ‘learning to remember’ in which subjects became better at employing more reliable past observations to determine the hidden state of the world.

## Introduction

Natural environments are replete with statistical structure and regularities over many spatial and temporal scales. Humans and other animals are adept at extracting this structure by building cognitive maps (Tolman, 1948) or world models (Daw et al., 2005; Gläscher et al., 2010), which support predictions of future states and requirements. This information can then be used to enable more efficient and effective actions and decisions, for instance allowing faster reactions to probable events (Niemi and Näätänen, 1981).

One critical aspect of prediction in environments involving temporal regularities is that it typically depends on memory, with the immediate sensory evidence alone being insufficient (Kaelbling et al., 1998). Such cases involve what is known as partial observability, as in a hidden Markov model (HMM), and pose difficulties for using a world model even when it has been learned. To achieve good performance, subjects must remember and integrate evidence provided by past observations. This demands the effective and adaptive use of forms of working memory (Zilli and Hasselmo, 2008; O’Reilly and Frank, 2006; Todd et al., 2009).

Hidden structures exist over a variety of timescales. Short times of a few seconds are associated with accumulate-to-bound decision making (Ratcliff and Rouder, 1998; Gold and Shadlen, 2002) or persistent activity states (Miyashita, 1988; Fuster, 1997; Frank et al., 2001). Very long times, perhaps even across days, are associated with macro-states or contexts (Haruno et al., 2001; Gershman et al., 2010). By contrast we consider a task in which the critical structure (which supervenes over shorter-time task requirements) typically concerns an intermediate scale of tens of seconds.

By analysing singular aspects of the behavioural data in the task we find that rats learn to use such medium-term structure to predict oncoming states and adjust their actions accordingly. However, their behaviour reveals imperfections which, when they arise, result from chance recent observations that are misleading as to the identity of the hidden state. We show how to account for their performance by assuming that they perform inference with an imperfect memory in an HMM that characterizes the environment. We also show that the subjects perform more competently as training progresses; and characterize this in terms of ‘learning to remember’, in which subjects decrease reliance on recent observations in favour of an increasingly dependable memory.

## Results

### Task and Experiment

We consider a cumulative handling time task (Breton et al., 2009; Solomon et al., 2017) in which rats hold down a lever for an experimenter-defined time period, called the price (*P*), in return for rewarding electrical stimulation of the medial forebrain bundle (Olds and Milner, 1954) at a fixed current and a given pulse frequency (*f*). In this paradigm, subjects experience many trials, each of which consists of an epoch during which price and frequency are fixed. Subjects may achieve the price cumulatively, over multiple presses during the trial. The duration of a trial (*D*) is 25 times the price (except for a minority of trials with price less than 1 second which last 25 seconds) allowing for many rewards to be obtained. This duration excludes a short, typically two second period following each reward termed the ‘black-out delay’ which allows for reward consumption and during which the lever is retracted and re-extended. Together frequency, price and duration define the experimentally-set parameters of a given trial. Since duration depends directly on price, being directly proportional to it in many trials, we follow convention (Breton et al., 2013; Niyogi et al., 2014a; Solomon et al., 2017) and use frequency and price but not duration in our models; see discussion.

At the beginning of each trial, a high frequency stimulation train, called a prime, is delivered. The subjects are then free to choose whether and when to engage with the lever. We analyse two major dependent variables. The first is the engagement probability (EP), which is the probability that subjects engage with the lever at all. The second, if subjects do indeed engage, is the initial response time (IRT), which is the time it takes them to first press the lever following the prime; we define these in more detail in materials and methods.

Trials come in a predictable cyclic triad consisting of ‘lead’, ‘test’ and ‘trail’ trial types (Fig 1A) separated by a fixed intertrial interval of 10s. Each trial type is associated with different frequencies, prices and durations (Fig 1B; shown with log base 10 here and subsequently), but are otherwise identical. When subjects know the frequency and price associated with a trial, and hence the worth of work, they typically choose an appropriate level of engagement with the lever. This is illustrated by the ethograms in Fig 1C in which the lever presses of a trained subject are plotted for different trial types (we ignore the post-reward ‘black-out delay’).

**Fig 1.**
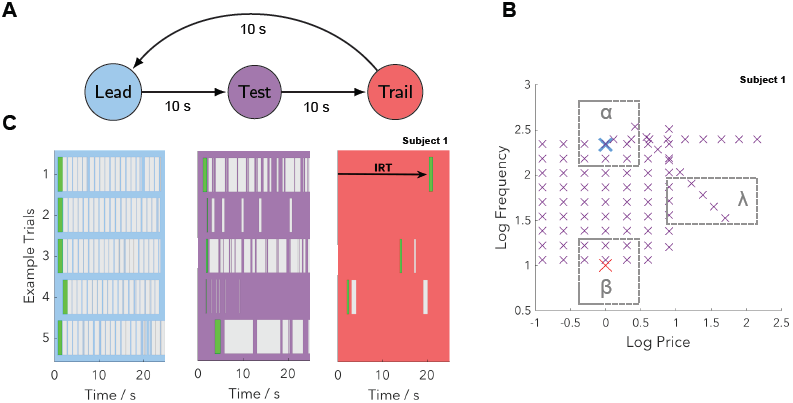
The structure of the experiment. (**A**) Trials come in a predictable cyclic triad. Each trial corresponds to a period of time where price and frequency are fixed. The intertrial interval is 10s. (**B**) Frequencies and prices associated with each trial type (for subject 1). Lead trials are highly rewarding with a fixed high pulse frequency and a short price (blue cross). Trail trials are negligibly rewarding with a fixed low pulse frequency and a short price (red cross). Test trails vary in frequency and price from trial to trial and so are variably rewarding (purple crosses). In addition to the crosses we also define regions α, β and λ (dashed grey rectangles) which are relevant for Fig 3. Note that in regions α and β, test trials are similar to lead and trail trials respectively, whereas in region λ test trials are dissimilar to both. (**C**) Responses from five example triads of trials from a trained subject. Grey bars correspond to the lever being depressed, with initial responses highlighted in green. Pressing is almost continuous on lead trials, varies on test trials from trial to trial (only the first 25s is shown) and is rare on trail trials. We label the IRT, which reflects subjects’ beliefs about the rewarding nature of the current trial before they have experienced any within-trial evidence.

For lead trials, which correspond to fixed, high-frequency stimulation with a short price of 1 second, subjects typically work the entire duration of the trial, as the high-frequency stimulation is highly rewarding. By contrast for trail trials, which have fixed, low-frequency stimulation at the same short price of 1 second, subjects barely work. Test trials, which involve a range of frequencies and prices which change from trial to trial (but are fixed across a particular trial) give rise to variable amounts of work, depending on the particular values of the frequency and price.

The data in the present paper are drawn from (Solomon et al., 2017) which describes in detail all aspects of the experiment, including training prior to the full task. Training involved a shaping protocol which eventually introduced lead, test and trail trials, enabling subjects to learn the cyclic triad structure. It used a more limited range of test frequencies and prices than was ultimately employed in the main experiment (Fig S1).

We studied a total of six subjects, each of which had experienced approximately 1500 triads of trials over a period of weeks. To allow adjustment from training to the full task we excluded the first 126 triads from our analysis, corresponding to one complete survey of the test trial frequencies and prices as defined in (Solomon et al., 2017). The number of surveys analysed for subjects 1-6 was 12, 10, 11, 8, 12, and 13 respectively, with each survey being acquired over 2 daily sessions, lasting approximately 6 to 7 hours each. Subjects 1-6 in this paper correspond to subjects F03, F09, F12, F16, F17 and F18 respectively in (Solomon et al., 2017). Whilst in general the results we describe apply to all six subjects, for simplicity we often display results in full for only subject 1, describing the remaining subjects using summary statistics. We report significance for individual subjects at the P < 0.05 level, with further details on exact p-values and of our methodology being referred to materials and methods.

### Subjects learn the task transition structure

Previous analysis of these data has primarily focused on behaviour during test trials, and in particular on responses occurring after the initial responses (Niyogi et al., 2014a; Niyogi et al., 2014b; Solomon et al., 2015; Solomon et al., 2017). Following Breton, 2013, we instead considered all trial types, and primarily focused on initial responses, since they reflect the subjects’ beliefs about the likely worth of a trial before they encounter any within-trial information. They are thus the best source of information about the subjects’ understanding of the cyclic triad structure. We characterized the initial responses by EPs and IRTs.

Fig 2 contrasts the performance of subjects when they have just begun training in the triad structure with the performance of the same subjects after they have been trained. For this analysis we exclude the first 5 triads during training as subjects were not always engaged in the new task when it first began but quickly learned to be. Analysis of the subsequent 20 triads for each subject revealed this, with EPs close to 1 and short IRTs for all trial types. These rapid and reliable responses likely reflected the subjects’ lack of understanding of the task structure, as if they predicted engagement with the lever to be valuable, or at least worth exploring, on all trial types. This was further supported by the finding that 5/6 subjects did not respond with a median trail trial IRT which was significantly longer than the median IRT of a combined distribution of lead and test trials (permutation test; *h*_1_). For the significant subject, the median trail trial IRT was not large (3.15s) and the EP was 1, and so this likely reflected initial stages of learning.

**Fig 2.**
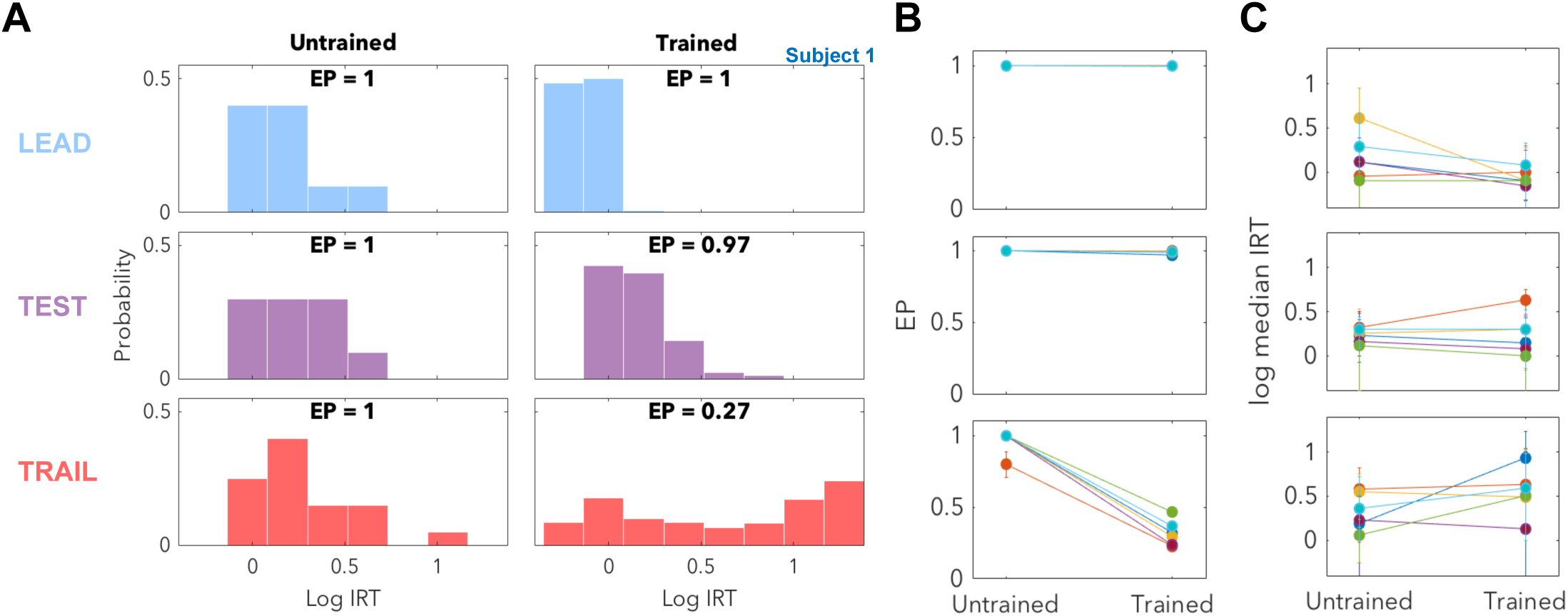
Subjects learn to predict oncoming trials. (**A**) We compare responses when subjects have just begun training with the triad structure to responses once trained. Early in training (left) the subject responds with short IRTs for all three trial types and EPs of 1, reflecting engagement in the task but an inability to predict the oncoming trial type. After training (right), IRTs reflect accurate prediction of oncoming lead and test trials, with certain engagement and rapid but generally distinguishable responses on the two trial types. For negligibly rewarding trail trials, the subject responds appropriately in the majority of cases, as indicated by both a low EP and a number of responses with long IRTs. However, in a minority of cases subjects also responded with short IRTs, which indicates inaccurate prediction of the trail trial. (**B**) For lead and test trials, EPs remained close to 1 (subject 1’s response in dark blue). On trail trials, EPs were found to decrease consistently for all 6 subjects (binomial proportion test; *h*_3_). (**C**) For lead trials, median IRTs remained short, and for 4/6 subjects became even shorter once trained (permutation test; *h*_4_), as subjects learned to predict the highly rewarding lead trial. For test trials, with their lower expected rewards, median IRTs remained relatively constant and were longer in trained subjects than lead trial IRTs for all subjects (permutation test; *h*_2_). For the poorly rewarding trail trials, median IRTs appeared not to change consistently, but we examine the properties of the trail trial distribution in more detail in Fig 3.

After the training period, the same subjects emitted very different initial responses for the different trial types. To a first approximation, the difference in these initial responses for trained subjects reflected the expected worth of the trial: the larger this worth, the greater the EP (up to a maximum of 1) and the shorter the IRT.

For lead and test trials, the EP was generally very close to 1, with test trial IRTs being slightly longer for all subjects than those for (the on average more valuable) lead trials (permutation test; *h*_2_). That test trial IRTs were longer than lead trial IRTs is interesting as this behaviour is seemingly suboptimal – subjects need to explore to find out the test trial’s value before they can determine the appropriate response, and waiting at the beginning of a trial reduces their potential to exploit the test trial if it is indeed of high value. We therefore interpret the longer latency on test trials as indicating a sub-optimal Pavlovian response to an accurate prediction of relatively lower expected future reward, an effect which has been observed elsewhere (Liu et al., 2000; Dayan et al., 2006). This type of response is convenient for our purposes as it indicates that subjects learned to accurately and differentially predict lead and test trials, even before they engaged with the lever.

Subjects responded very differently on the negligibly rewarding trail trials. EPs were typically small, and when subjects did engage, the resulting IRTs were often long. However, on a substantial fraction of occasions, the IRTs were instead short, which is surprising because trail trials were designed to be effectively worthless to the subject. We explore the possibility that the pattern of long and short IRTs is a signature of subjects’ inability to predict trail trials perfectly, and are thus a result of erroneous inference. They therefore provide a window into the subjects’ inferential processes.

### Misleading evidence leads to mistaken state inference

Trail trials are preceded by test trials, which involve a range of different frequencies and prices. Some of these conditions resemble either lead (region α, Fig 1B) or trail (region *β*, Fig 1B) trials. According to the task transition structure, lead or trail trials are followed by test or lead trials respectively, both of which are associated with high EPs and low IRTs in subjects’ initial responses. We therefore considered the possibility that short IRTs on trail trials arose when the subjects had been confused by the preceding test trial, but had applied their good knowledge of the transition structure (Fig 3A; see also Breton, 2013).

**Fig 3.**
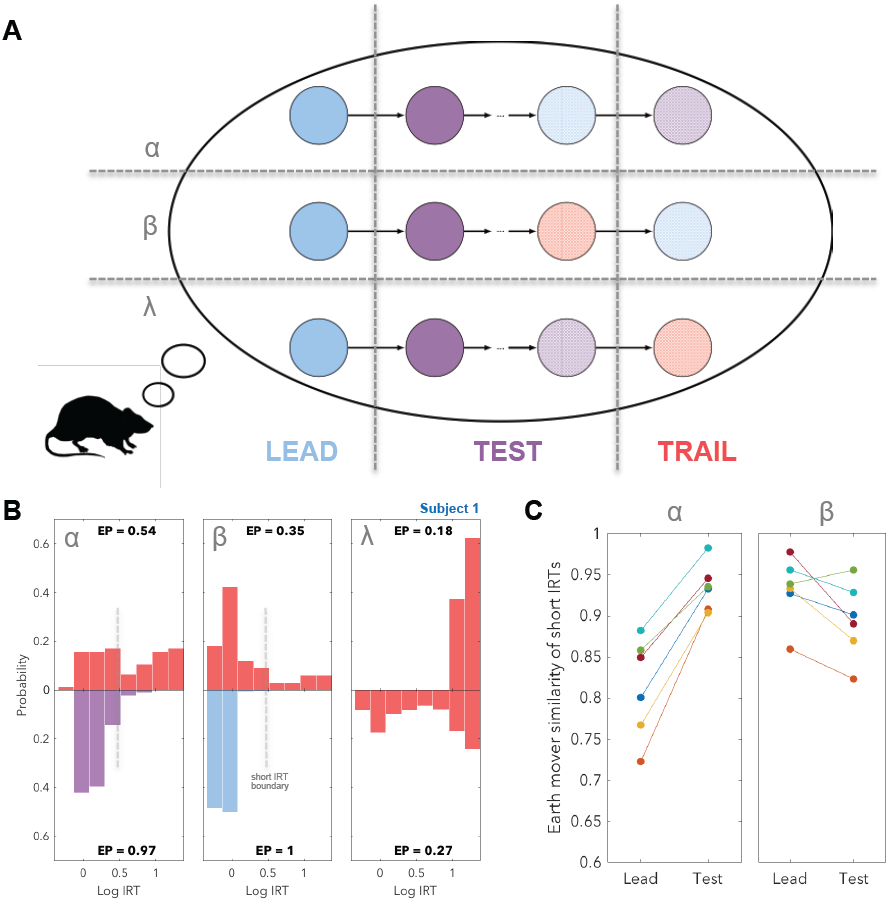
Short IRTs on comparatively worthless trail trials as mistaken inferences. (**A**) When a test trial had a frequency and price similar to that of either a lead trial or a trail trial (regions α and β respectively; Fig 1B), the subject’s belief about the trial type that they were experiencing may have been mistaken by the time the trial ended. If this incorrect belief was combined with a correct understanding of the transition structure of the task, then the subject would have expected the next trial to be a test or lead trial respectively, rather than a trail trial, and so would have chosen a short IRT rather than no response or a long IRT. By contrast, if the frequency and price were dissimilar to both lead and test trials (region λ; Fig 1B), then the test trial would be unambiguous and the subject would either not respond at all, or would elect a long IRT on what it considered to be the subsequent trail trial. These effects are likely to be probabilistic, which we indicate by lighter shading. (**B**) We tested this hypothesis by examining trail trial responses given that the preceding test trial’s frequency and price were in regions α, β or λ. To aid visual comparison of histograms we use a mirror plot. Left: When test trials had similar frequency and price to lead trials (region α), the resulting distribution of short trail trial IRTs (upper) was test-like (the lower left plot shows the actual distribution of IRTs on test trials). Middle: when test trials were similar to trail trials (region β) the resulting distribution of short trail trial IRTs (upper) was lead-like (lower plot). Right: when test trials were dissimilar to both lead and test trials (region λ), short IRTs were no longer observed (upper) despite these responses being common in the trail trial distribution which includes all preceding frequencies and prices (lower). (**C**) This confusion effect is trial type specific. Short IRTs on trail trials following test trials in region α are more similar to test trial IRTs than to lead trial IRTs for all 6 subjects (permutation test; *h*_5_). Similarly, short IRTs on trail trials following test trials in region β are more similar to lead trial IRTs than to test trial IRTs for 3/6 subjects with the difference not being significant for the remaining subjects (permutation test; *h*_6_).

To test this hypothesis, we sorted the trail trial IRTs by the frequency and price of the previous test trial (Fig 3B). Indeed, when the test trial had similar frequency and price to a lead trial (region α), the resulting distribution of short trail trial IRTs resembled that of a test trial. This is consistent with the subject inferring the test trial to be a lead trial and hence the subsequent trail trial to be a test trial. Likewise, we found that when the test trial was similar to a trail trial (region *β*) the resulting distribution of trail trial IRTs was similar to that of a lead trial, again consistent with expectations. For test trial frequency-price combinations dissimilar to those of either lead or trail trials (region λ, Fig 1B), subjects were rarely confused, and so short IRTs occurred much more rarely and EPs were much lower (Fig S2).

To quantify whether the short IRTs sorted in this way are more lead-like or trail-like respectively we calculated the earth mover’s similarity between these distributions and lead and trail distributions (Fig 3C). We define the earth mover’s similarity to be 1 minus the earth mover’s distance (or equivalently 1 minus the Wasserstein distance). To select only short IRTs, we eliminated IRTs greater than the 95th percentile of the test trial distribution. We found that for all 6 subjects, responses on a trail trial following a lead-like test trial were significantly more test-like than lead like (permutation test; *h*_5_). Similarly, responses on a trail trial following a trail-like test trial were significantly more lead-like than test-like for 3/6 subjects (permutation test; *h*_6_).

Having discovered this confusion effect, we investigated it in more detail by considering the separate influences of frequency, price and duration. We found that frequency strongly influenced subjects’ inferences (Fig 4A): for intermediate, and therefore not misleading, frequencies, subjects were much less likely to respond rapidly on a trail trial, even when price and duration were misleading. Similarly, we found subjects were sensitive to price and/or duration (Fig 4B), as when these were long, and therefore not misleading (since lead and trail trials ubiquitously had price of 1s), subjects were much less likely to respond rapidly even when the frequency was misleading. Finally, we also examined the minority of cases when price was not misleading but duration was (Fig 4C). These arose when price was less than 1 second, as the duration was fixed at 25 seconds rather than being 25 times the price, as otherwise. We show that subjects were sensitive to the price on the preceding test trial when its frequency was high but not when it was low. We speculate about the reason for this in the discussion, but do not seek to model it as its effect is subtle, only influencing a fraction of the data.

**Fig 4.**
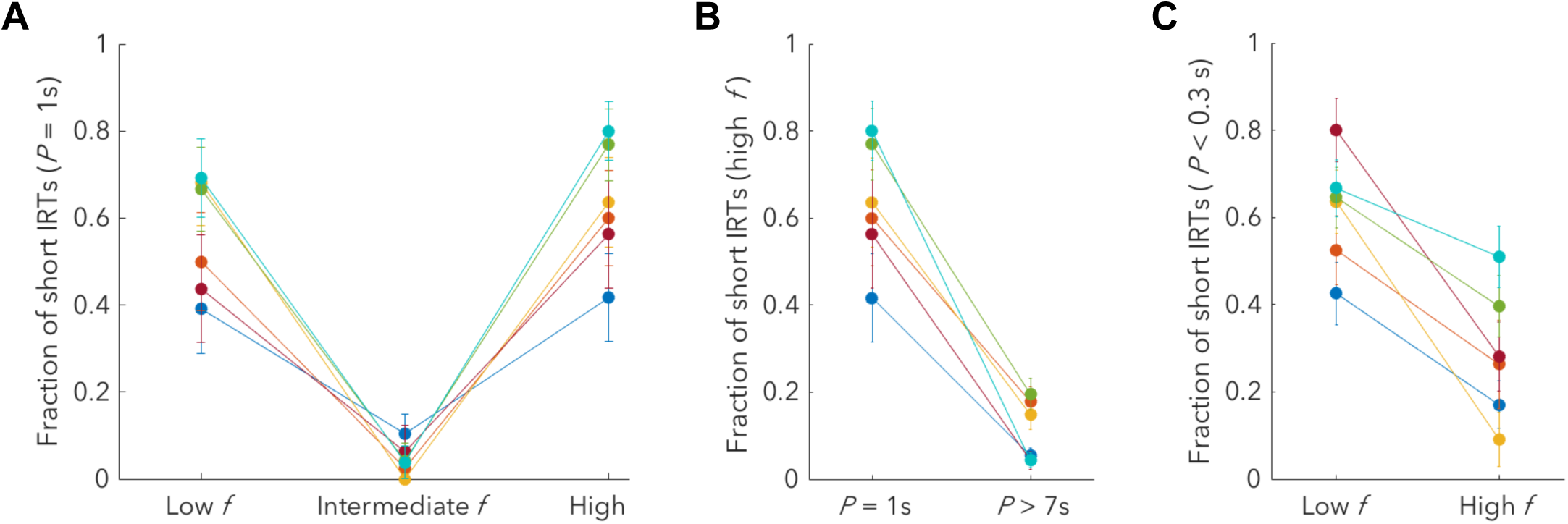
Subjects use multiple sources of evidence from the preceding test trial to determine a response on the trail trial. (**A**) Intermediate test trial frequencies only very rarely lead to short trail trial IRTs, even when price and duration are misleading. This indicates that subjects can use frequency to determine the appropriate response when this frequency is different from that of lead or trail trials; see materials and methods for a definition of these regions. This difference is significant for all subjects when comparing intermediate to both ‘Low f’ (binomial proportion test; *h*_7_) and ‘High f’ (binomial proportion test; *h*_8_) categories. (**B**) High test trial prices also only rarely lead to short trail trial IRTs even when frequency is misleading (here we show for high, lead-like frequency). As duration is perfectly correlated with price for prices of 1 second or greater, this indicates that subjects can use price and/or duration to determine the appropriate response. The difference between the two categories is significant for all subjects (binomial proportion test; *h*_9_). (**C**) When test trial price is short, test trial duration remains at 25 seconds, thus we consider cases in which price is not misleading (*<* 0.3s) but duration and frequency are. Short trail trial IRTs depend on the frequency of the preceding test trial. When the frequency is low, subjects respond with a similar fraction of short responses as for a price of 1s (Figure 4A; left), indicating price insensitivity. However, when the frequency is high, subjects are price sensitive, with a decreased fraction of short responses. This difference was significant for 4/6 subjects (binomial proportion test; *h*_10_).

Whilst the description of confusion outlined in Fig 3 provides a clear, model-agnostic, account of the varied responses on trail trials, it only provides a simplified, deterministic picture of this process. We therefore built a probabilistic model, incorporating our understanding from Fig 4A;B, in order to describe this more precisely.

### Modelling the inference process

The task itself can be described in the form of an HMM, with hidden states representing the trial type, and a binary transition matrix reflecting the deterministic cyclic triad structure. Given the predominant regularities in responses, highlighted earlier, we assume that subjects have learned this essential structure, associated with a task transition matrix (**A**), which acts on the subject’s belief state when transitioning between trials.

Fig 5 captures the key steps of correct and incorrect inference. First, we assume that at the end of a lead trial, the subjects were correctly certain that this was their current state. Matrix **A** operates on this belief such that the subject’s certainty would propagate into certainty that a test trial would come next. This assumption is based on the fact that EPs on test trials are near 1 for all subjects and IRTs are short (but different from lead trials), suggesting negligible confusion at this point.

**Fig 5.**
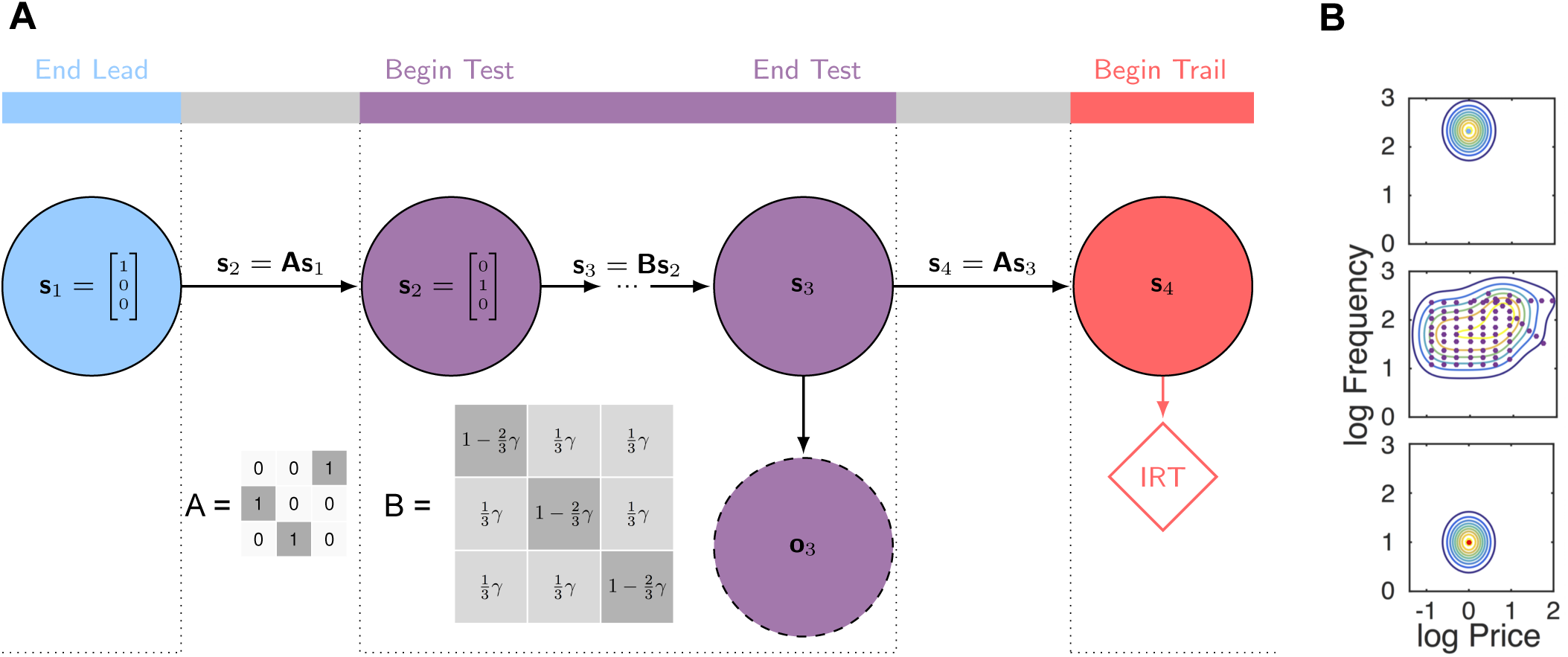
Modelling the inference process. (**A**) We characterize subjects as building an HMM generative model of the task and performing recognition to produce posterior subjective beliefs over the trial types. In our model, at the end of a lead trial the subject is certain it is on a lead trial (*s*_1_). As it has learned the transition structure, described by matrix **A**, it is therefore certain it is on a test trial at the beginning of a test trial (*s*_2_). If recognition was perfect, this knowledge would persist through the test trial; we model subjects’ imperfection as arising from forgetting during the test trial. For convenience, we describe forgetting with probability *γ* as arising from an incorrect generative model, described by transition matrix **B**. By the end of the test trial, the evidence from memory is integrated with the within-trial evidence provided by observations (*o*_3_) of frequency and price. This leads to a posterior belief (*s*_3_), which then leads to the subjective belief about the trial type at the beginning of what is actually the trail trial (*s*_4_). This can then be used to generate a response: either no engagement or engagement with an associated IRT. (**B**) We describe the association between points in frequency-price space and trial type using a mixture of Gaussians centered at the experimentally utilised points for lead (top), test (middle) and trail (bottom) trials. We introduce a standard deviation parameter (*σ*) which is shared across all points.

If the subjects’ inference was perfect, they would continue to believe that they were in a test trial throughout. However, this continued belief depends on a perfect working memory. We model imperfection as arising from an incorrect generative model (but see the discussion) involving an intermediate forgetting matrix (**B**), which allows for the possibility that subjects could forget the current trial type. Matrix **B** is parameterized by a scalar *γ*. If *γ* = 1, forgetting is complete, making all states *a priori* equally likely at the end of the test trial; if *γ* = 0, there is perfect memory, implying that the test trial would remain unambiguously known.

However, the subjects did not need to rely solely on their imperfect memories. They also received potentially noisy observations of the frequency and price during the test trial, and could integrate these into their beliefs. To determine the strength of evidence provided by these recent observations we introduce a standard deviation parameter (*σ*), using this to generate a mixture of Gaussians distribution (or equivalently kernel density estimate) in log frequency-price space associated with each trial. To specify the centre of each Gaussian, we use the real points in log frequency-price space experienced by each subject during the experiment, scaling the mixture weights in proportion to the number of times with which they were utilized.

For a given triad of trials, probabilistic integration according to the HMM can be described using Bayes rule as:

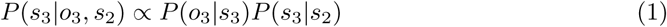

where *s*_3_ is the inferred trial type at the end of a test trial, *o*_3_ is the observed frequency and price (*f, P*) on the test trial and *s*_2_ is the state at the beginning of the test trial.

This makes clear the influence of both recent observations, *P* (*o*_3_*|s*_3_), and slonger-term evidence from memory, *P* (*s*_3_ *s*_2_), on the posterior belief at the end of a test trial. Having determined this belief we find the belief at the beginning of a trail trial, *P* (*s*_4_ *o*_3_, *s*_2_), simply by applying the task transition matrix **A**.

We then calculate the probability of a particular response according to:

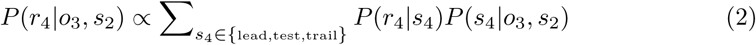

where *r*_4_ is the response at the beginning of a trail trial (including no responses) and the summation is over the three possible trial types.

To calculate the probability of IRTs given a known trial type we used non-parametric fitting of lead, test and non-confusing trail trial IRTs (Fig S3). The latter distribution was found by only selecting trail trials which followed test trials in region λ, which thus largely eliminated short IRTs. In order to fit parameters *γ* and *σ* we used the real responses generated by the subjects and maximized the sum of the log likelihoods of those responses with respect to the parameters.

Having built the HMM we then split the data into three tertiles (details outlined in the following subsection), and determined the maximum likelihood estimate (MLE) of the parameters independently for each tertile. We were then able to simulate response distributions by sampling from 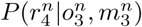 where *n* indexes a particular triad of trials. We found that we were able to recover the pattern of short and long IRTs present in the real data, closely matching the observed distribution of EPs and IRTs for 5/6 subjects (Fig 6A;B;C). When sorting the simulated data by the previous test frequency and price in the same manner as before, the simulated data was found to match the real data well (Fig 6D), indicating that the model is able to account for the observed confusion.

**Fig 6.**
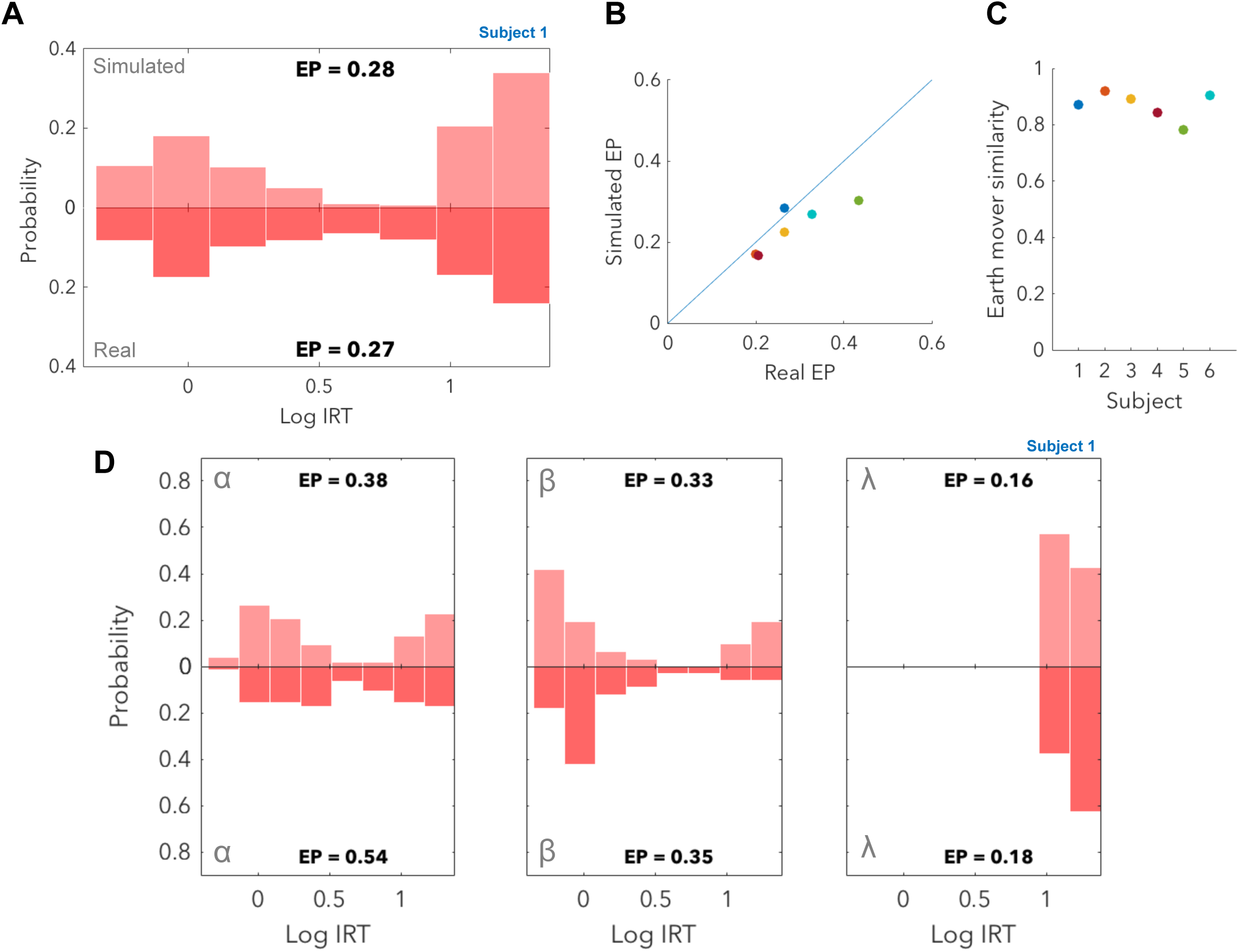
Simulated responses capture the process of mistaken inference. (**A**) By fitting model parameters and simulating responses (upper), we are able to recover the distribution of short and long IRTs observed in the data (lower). (**B**) Simulated EPs are similar to real EPs except for subject 5 (green). (**C**) The simulated distributions of IRTs have earth mover’s similarities to the real distribution above 0.8 except for subject 5. (**D**) By sorting responses into regions as in Fig 3B. we find that simulated distributions are similar to the real distributions indicating that our model is able to capture the confusion effect.

To investigate simpler versions of the model could provide a more parsimonious explanation for the observed responses we also tested models in which subjects only used evidence from one of frequency or price but not both, and also a model which used frequency and price but did not utilize a *γ* parameter (Table 1). These gave much poorer fits however as reflected by higher BIC scores, justifying the full version of the model over these alternatives.

**Table 1.** The increase in the BIC score for simpler versions of the model.

| Subject | No $\gamma$ | No $f$ | No $P$ |
| --- | --- | --- | --- |
| 1 | 208.6 | 15.2 | 30.4 |
| 2 | 33.8 | 26.8 | 33.1 |
| 3 | 33.0 | 9.6 | 6.1 |
| 4 | 126.6 | 30.0 | 20.9 |
| 5 | 63.4 | 28.6 | 76.3 |
| 6 | 69.4 | 47.8 | 70.0 |

### Inference improves with experience

Subjects typically encountered well over a thousand triads of trials. We therefore analysed improvements on the task with experience by dividing the data by triads into three sequential tertiles. When comparing the final tertile to the first tertile for subject 1 we observe a marked decrease both in the EP and in the probability of short IRTs on trail trials (Fig 7A). To analyse this across all subjects we calculated the fraction of short IRTs for each tertile and found it to be significantly decreased for 4/6 subjects, with the remaining subjects showing no significant change (Fig 7B; permutation test; *h*_11_). Taken together, this indicates that by the final tertile most subjects had improved their ability to track their progress through the task, as even on misleading trials they were rarely confused.

**Fig 7.**
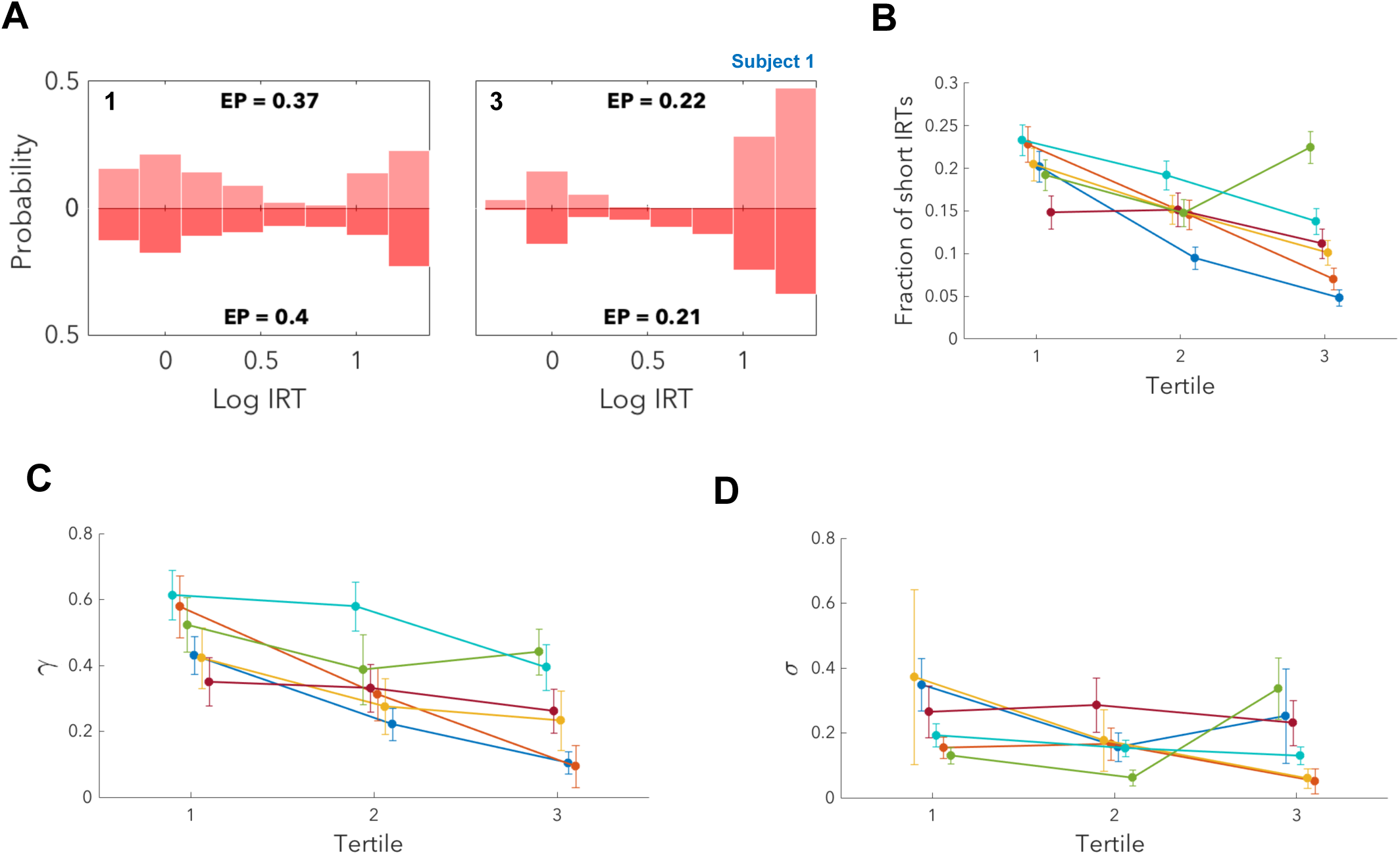
Mistaken inference becomes less likely with experience, as subjects learn to remember. (**A**) Upper: simulated; lower: real. We divided the data into three tertiles, fit parameters independently for each tertile and simulated responses. We illustrate the first and last tertile in which the subject both lowers its EP and also decreases the probability of short IRTs in cases when it does respond. (**B**) By plotting the fraction of short IRTs in each tertile we find a significant decrease from first to last tertile for 4/6 subjects (permutation test; *h*_11_) indicating that these subjects improve in their ability to identify the trail trial. The remaining two subjects shown no significant change. (**C**) By calculating *γ* for each tertile we find that 4/6 subjects show a significant decrease in the MLE estimate of *γ* from the first to last tertiles, with the remaining two subjects showing no significant change (permutation test; *h*_12_). Although significance was tested using a permutation test, we illustrate errorbars using the mean square error in the MLE of the parameters. The decrease in *γ* over time for the majority of subjects suggests a process by which subjects learn to remember. (**D**) There is no significant change in the MLE of *σ* for 3/6 subjects with two subjects showing a significant decrease and one a significant increase (permutation test; *h*_13_). We therefore do not find strong evidence to suggest that improvements in performance in the majority of subjects was due to a more accurate association of frequency and price with the trial type.

In order to understand these changes in the context of our model, we fit model parameters independently to each tertile. MLEs of the parameters identified significantly lower values of *γ*, the probability of forgetting, in the last tertile relative to the first tertile for 4/6 subjects, with the remaining subjects not showing a significant change (Fig 7C; permutation test; *h*_12_). The subjects for which this parameter changed significantly corresponded to those which had shown a significant decrease in the fraction of short IRTs. This suggests that over time, the majority of the subjects learned to remember the previous trial type better and so improved their identification of the test and subsequent trail trials.

We also examined changes in the MLEs of the parameter *σ* across tertiles and found no significant change for 3/6 subjects, a significant decrease for two subjects and a significant increase for one subject (Fig 7D; permutation test; *h*_13_). This indicates that for most subjects there is no evidence that improvements in performance can be attributed to a more accurate association of frequency and price with the appropriate trial type.

Finally, to assess the linear correlation between estimates of the parameters *γ* and *σ* we calculated the Pearson correlation coefficient from the negative inverse Hessian evaluated at the MLE (Fig S4; see materials and methods for further details). We determined this coefficient separately for each subject and for each tertile, and typically found a negative value between −0.3 and −0.7, indicating moderate anticorrelation in the estimated parameters.

## Discussion

We have shown that subjects learned a model of the world which reflected an experimentally defined transition structure. However, we also identified a small fraction of trials where behaviour seemingly went awry, as evidenced by subjects responding rapidly in advance of unrewarding trials. We demonstrated that these responses could be attributed to mistaken inference of the trial type, and described this process using an HMM. This involved introducing two parameters: *σ*, which influenced the mapping from observations during the trial to the inferred trial type; and *γ*, which represented the fidelity of subjects’ memory of the previous state. We observed that the latter parameter decreased significantly over the course of hundreds of triads for the majority of subjects, suggesting a process by which subjects learn to remember.

An important part of the work we have described is not only demonstrating subjects’ abilities to learn structure in their environment but also in building a statistical model which describes inference in this context. The model developed was clearly defined and involved parameters which were interpretable, allowing for greater insight into the changing role of both memory and recent observations.

The representation used in our model proposed that subjects maintain belief states in an HMM. It is also conceivable that they might instead have adopted a less compressed, history-based representation of state, by storing the frequency and price of previous trials (either explicitly or implicitly). It is hard to distinguish these based only on behaviour (particularly given the relative paucity of errors); but this would, of course, still constitute a functional form of world model.

In our preferred representation, observed ‘mistakes’, corresponding to short IRTs on worthless trail trials, are due to mistaken inference of the hidden state. To support this claim we demonstrated that when the inference problem was easy, such as following lead, trail or non-confusing test trials, subjects’ responses reflected clear understanding of the structure. By contrast, when inference was hard, subjects more frequently responded inappropriately in a way which we were able to predict. Although we couched the problem through a forgetting matrix in the generative model, this really captures a form of approximate recognition. It is interesting to consider the alternative possibility that this incorrect generative model actually constituted the subjects’ subjective beliefs about the environment. This would allow transitions between states to occur at any point during a trial as a result of misleading observations. In this interpretation, observed improvements in performance over time could be attributed to subjects learning a better model, rather than improving their inference. This could give rise to exactly the model we used, with *γ* determining the model’s probability of switching belief to another state instead of the probability of forgetting. One trouble with this interpretation is that subjects always experienced each trial as being deterministically stable across time, with no change in either price or frequency. We suggest that subjects’ world models would incorporate this regularity.

Our model is starkly simple, using only two parameters to predict behaviour without reference to the detailed microstructure of a given trial, such as the number of reward encounters or the average reward rate. Fig 4C provides a hint that the former factor can be influential, as increased sensitivity to price following high frequency test trials may have resulted from an increased number of lever presses on these trials and thus a more accurate perception of price.

Nevertheless, we made this choice to capture and highlight the predominant effects observed across subjects whilst also maintaining interpretability. In turn, this allowed us to identify a significant change in the *γ* parameter in the majority of subjects, a finding supported both by calculating the standard error in the mean of these parameters and by permutation testing. In addition to identifying parameters we also determined linear correlations between them at the MLE, and found that estimates of *γ* and *σ* were moderately anticorrelated. This finding can be understood intuitively by considering a variation in the parameters such that *γ* is increased but *σ* is decreased. In this case, the probability of forgetting increases but frequency and price are now more accurately perceived, resulting in a ‘trade-off’ between the two parameters when predicting behavioural performance. However, this trade-off is only partial, ultimately allowing for separate estimation of *γ* and *σ* with sufficient data.

Another related aspect of fitting our model was the choice not to use trial duration in addition to price to predict responses. As alluded to earlier, this was due to the strong correlation between duration and price, which implied that using either would produce similar results. On the other hand, we were able to show that subjects do use both frequency and price/duration, indicating that they successfully combined multiple sources of evidence in the inferential process.

Two aspects of learning merit future work. One is how the subjects learned the overall model of the world over early training – particularly given their initially imperfect memories and their ignorance of the number of potential states. One promising approach is to consider a non-parametric statistical structure such as an infinite hidden Markov model (Beal et al., 2002). The second aspect of learning concerns learning to remember. The task demands in excess of 45 seconds of memory in order for subjects to utilise information from two trials back. However, limits on the subjects’ capacities, and the relationship between their willingness to deploy this expensive resource and the resulting distribution of rewards (Kurzban et al., 2013; Botvinick and Braver, 2015) are unclear. Unfortunately, the structure of the task made assessing the dynamics of memory within a trial difficult to uncover; whilst longer trials might be expected to result in more forgetting and decreased accuracy in our task, this effect is confounded by the ability of subjects to use an extended price/duration to infer trial type more accurately.

Finally, understanding the neural underpinnings of the diverse processes involved in this task provides an exciting challenge for future research. In the case of working memory, its functioning is thought to be supported by persistent activity in a number of brain regions, including medial prefrontal cortex (Wang and Cai, 2006; Yoon et al., 2008; Horst and Laubach, 2009; Yang et al., 2014), entorinhal cortex (Hölscher and Schmidt, 1994; Egorov et al., 2002) and the hippocampus (Wang and Cai, 2006; Yoon et al., 2008). Measurements in these regions would potentially illuminate the precise role of working memory in this task, perhaps allowing for potential neural correlates of learning to remember to be uncovered. For evidence of neural representations of task structure, the hippocampus provides a natural candidate (Constantinescu et al., 2016; Garvert et al., 2017), and orbitofrontal cortex might similarly be suitable, given implications that it can encode a probability distribution over hidden causes (Wilson et al., 2014; Gershman et al., 2015; Schuck et al., 2016; Chan et al., 2016).

## Materials and methods

For a full description of the experimental methodology see Solomon et al., 2017. We outline here elements of the modelling methodology.

### Analysis

Since all trial types terminate after set intervals (25s for lead and trail; a variable duration for the test), some care is necessary with the resulting censoring of the time during which the subjects could engage. Furthermore, we occasionally observed cases towards the end of the trail trial in which the subject briefly pressed the lever for such a short time that there was no possibility of obtaining reward. This might have been a Pavlovian reaction to the expectation of the upcoming lead trial.

To avoid problems from these cases, we counted a trial as having been engaged in for the purposes of the EP if at least one reward was obtained, and we only considered IRTs (defined as the time taken from the beginning of a trial to press the lever for the first time) on those same trials. This constraint implies that initial responses after 24s on lead and trail trials would be impossible as there remains insufficient time to obtain a reward; so we only examine the properties of the IRT below this value. For test trials, which have variable trial duration, this ignored potential IRTs much larger than 24 seconds. However, in practice such cases were extremely rare (Fig 2A).

One facet of the experimental design is that the subjects received idiosyncratic calibrated frequencies of brain stimulation reward. We duly defined lower (*l*) and upper (*u*) boundaries of the regions *α, β* and *λ* separately for each animal; these also defined the boundaries dividing low, intermediate and high frequencies in Figure 4. Table 2 summarises these values. All frequencies are in Hertz and all prices are in seconds.

**Table 2.** Frequencies and prices used for lead and trail trials and for boundaries of regions *α β* and *λ*.

| Subject | $f_{\text{lead}}$ | $P_{\text{lead}}$ | $f_{\text{trail}}$ | $P_{\text{trail}}$ | $f_u^\beta$ | $f_l^\alpha$ | $P_l^{\beta/\alpha}$ | $P_u^{\beta/\alpha}$ | $f_l^\lambda$ | $f_u^\lambda$ | $P_l^\lambda$ |
| --- | --- | --- | --- | --- | --- | --- | --- | --- | --- | --- | --- |
| 1 | 217.4 | 1.0 | 10.0 | 1.0 | 20.0 | 125.9 | 0.4 | 3.9 | 31.6 | 79.4 | 8.1 |
| 2 | 196.1 | 1.0 | 10.0 | 1.0 | 35.5 | 100.0 | 0.4 | 3.9 | 63.1 | 125.9 | 4.1 |
| 3 | 250.0 | 1.0 | 10.0 | 1.0 | 44.7 | 158.5 | 0.4 | 3.9 | 79.4 | 158.5 | 4.1 |
| 4 | 200.0 | 1.0 | 10.0 | 1.0 | 31.6 | 100.0 | 0.4 | 3.9 | 50.1 | 79.4 | 4.1 |
| 5 | 250.0 | 1.0 | 10.0 | 1.0 | 28.2 | 125.9 | 0.4 | 3.9 | 39.8 | 100.0 | 4.1 |
| 6 | 163.9 | 1.0 | 10.0 | 1.0 | 25.1 | 79.4 | 0.4 | 3.9 | 39.8 | 100.0 | 4.1 |

### Statistical tests

We tested for statistical significance using two-tailed permutation and binomial proportion tests. Permutation tests were used to determine the probability that the observed difference in the test statistics between classes would occur for class labels which were randomly permuted. In all cases we used 1000 simulations.

One non-trivial usage of the permutation test was to see if changes in the MLE of model parameters was significant. For this we determined the MLE of the model parameters in the first and last tertiles for data in which the time labels were permuted randomly and calculated the absolute difference between these parameter values. This was repeated 1000 times in order to generate a distribution of differences. We then tested if the absolute difference in the MLE of the parameters for the non-permuted data was significant (greater than the 95th percentile).

The binomial proportion tests were used to determine the probability of the equality of two binomial proportions for two observed distributions. To compute this we evaluated the test statistic:

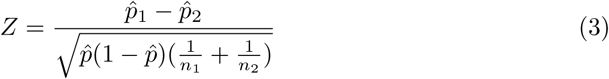

where 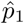 and 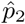 are the empirical probabilities, *n*_1_ and *n*_2_ the corresponding number of observations and:

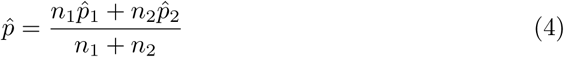

We then calculated p-values from *Z* using the normal approximation.

### Null hypotheses and p-values

Our null hypotheses referenced in the Results section were as follows:

*h*_1_: Trail trial IRTs have the same median as a combined grouping of lead and test IRTs for untrained subjects (permutation test)

*h*_2_: Test trial IRTs have the same median as lead trial IRTs for trained subjects (permutation test)

*h*_3_: EPs on trail trials are the same for trained subjects as they are for untrained subjects (binomial proportion test)

*h*_4_: Lead trial IRTs have the same median for trained subjects and untrained subjects (permutation test)

*h*_5_: Test trial IRTs are equally similar to trail trial IRTs with preceding test trials in region *α* as lead trial IRTs (permutation test)

*h*_6_: Lead trial IRTs are equally similar to trail trial IRTs with preceding test trials in region *β* as test trial IRTs (permutation test)

*h*_7_: The fraction of short trail trial IRTs is the same for the ‘Intermediate’ category as for the ‘Low f’ category, with P = 1s (binomial proportion test)

*h*_8_: The fraction of short trail trial IRTs is the same for the ‘Intermediate’ category as for the ‘High f’ category, with P = 1s (binomial proportion test)

*h*_9_: The fraction of short trail trial IRTs is the same for the ‘P = 1s’ category as for the ‘P > 7s’ category, with high f (binomial proportion test)

*h*_10_: The fraction of short trail trial IRTs is the same for the ‘Low f’ category as for the ‘High f’ category, with P < 0.3s (binomial proportion test)

*h*_11_: The fraction of short trail trial IRTs is the same in the final tertile as it is in the first tertile (binomial proportion test)

*h*_12_: The MLE of *γ* is the same in the final tertile as it is in the first tertile (permutation test)

*h*_13_: The MLE of *σ* is the same in the final tertile as it is in the first tertile (permutation test)

The P-values for these hypotheses for all subjects are listed in Table 3.

**Table 3.** P-values for null hypotheses.

| Null hypothesis | Subject 1 | Subject 2 | Subject 3 | Subject 4 | Subject 5 | Subject 6 |
| --- | --- | --- | --- | --- | --- | --- |
| $h_1$ | 0.506 | 0.001 | 0.472 | 0.200 | 0.488 | 0.543 |
| $h_2$ | < 0.001 | < 0.001 | < 0.001 | < 0.001 | < 0.001 | < 0.001 |
| $h_3$ | < 0.001 | < 0.001 | < 0.001 | < 0.001 | < 0.001 | < 0.001 |
| $h_4$ | 0.024 | < 0.001 | 0.532 | 0.025 | < 0.001 | 0.723 |
| $h_5$ | < 0.001 | < 0.001 | < 0.001 | 0.003 | 0.003 | 0.001 |
| $h_6$ | 0.023 | 0.181 | < 0.001 | 0.003 | 0.414 | 0.177 |
| $h_7$ | 0.004 | < 0.001 | < 0.001 | 0.014 | < 0.001 | < 0.001 |
| $h_8$ | 0.002 | < 0.001 | < 0.001 | 0.002 | < 0.001 | < 0.001 |
| $h_9$ | < 0.001 | < 0.001 | < 0.001 | < 0.001 | < 0.001 | < 0.001 |
| $h_{10}$ | 0.007 | 0.058 | < 0.001 | < 0.001 | 0.014 | 0.108 |
| $h_{11}$ | < 0.001 | < 0.001 | < 0.001 | 0.165 | 0.210 | < 0.001 |
| $h_{12}$ | < 0.001 | < 0.001 | 0.049 | 0.204 | 0.232 | < 0.001 |
| $h_{13}$ | 0.355 | 0.044 | 0.065 | 0.743 | < 0.001 | 0.022 |

### Model Comparison

We calculate the Bayesian Information Criterion (BIC) for a given model M according to:

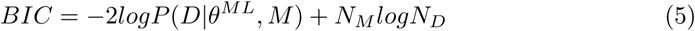

Where *D* is observed data, *θ*^*ML*^ are the maximum likelihood parameters of the model, *N*_*M*_ is the number of model parameters and *N*_*D*_ is the number of data points.

As we split the data into tertiles, we calculate the BIC for each tertile first and sum these to form an overall BIC for each subject.

### Comparison of model parameters across tertiles

When comparing model parameters across tertiles, for illustration in Fig 7C;D, we determined the standard error in the MLE of the parameters *θ* = (*γ, σ*) according to:

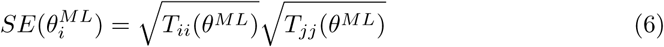

where 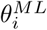 is the MLE of the parameter in question, *T*_*ii*_ is the i^th^ diagonal element of the matrix *T* = −H^−1^, the negative inverse of the Hessian *H*, defined as:

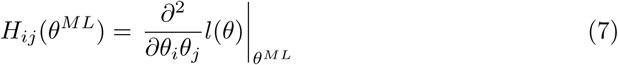

where *l*(*θ*) is the log likelihood.

As the matrix *T* is an estimator of the asymptotic covariance matrix we use it to determine the Pearson correlation coefficient:

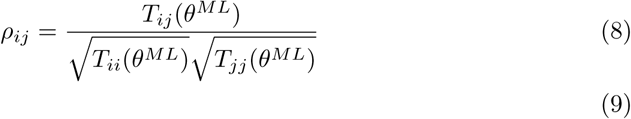

## Supporting information

**Fig S1.**
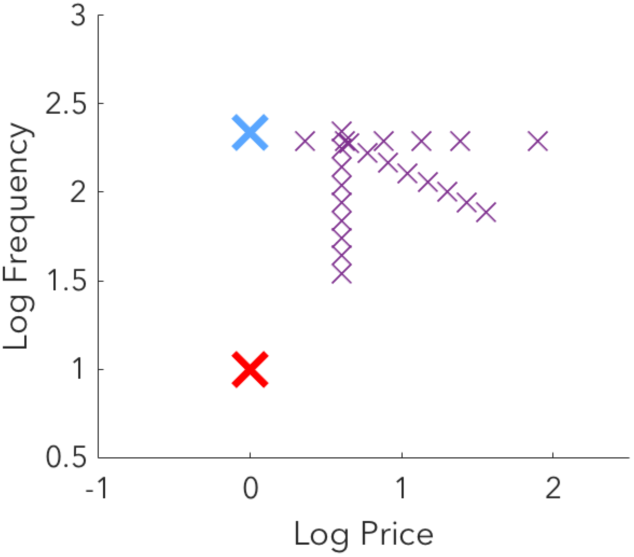
Training with the triad structure involved a more limited range of frequencies and prices. Frequencies and prices used during training for subject 1 are shown. The range of frequencies and prices is more limited than that employed in the full experiment.

**Fig S2.**
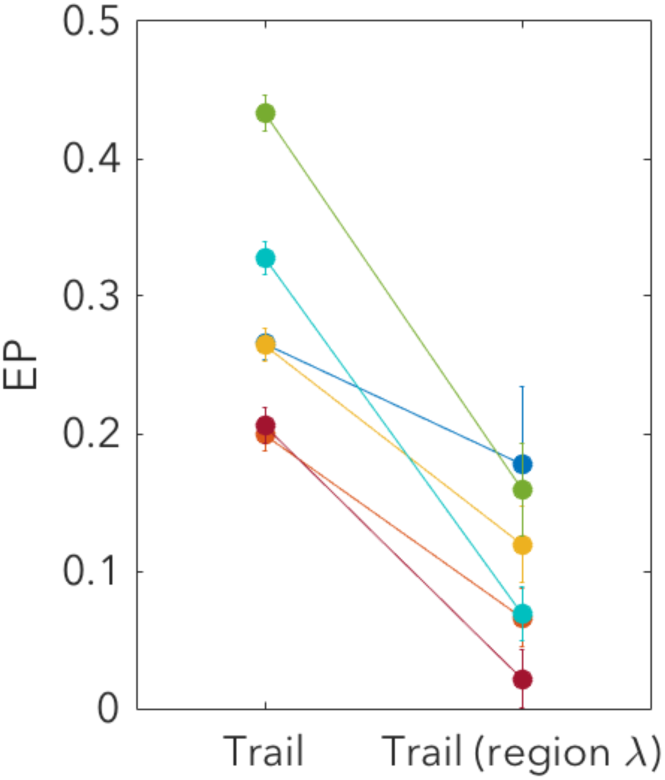
EP for trail trials is reduced when preceding test trial is in region λ. When trail trial responses are filtered such that only those with preceding test trials in region λ are included, a decrease in the EP is observed

**Fig S3.**
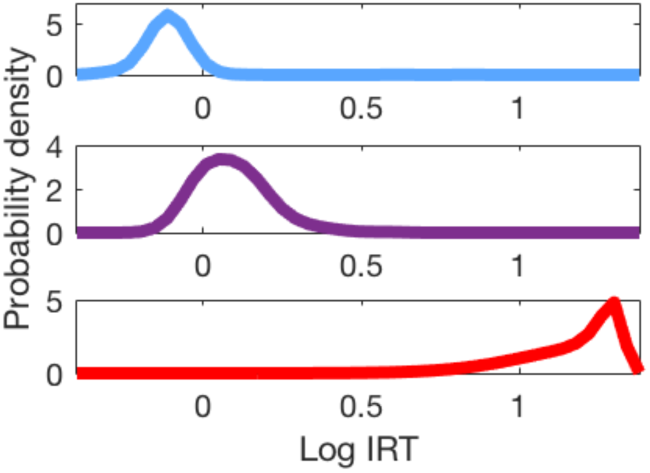
Determining the likelihood of responses given a posterior state. We evaluated the probability of the observed responses given certainty about the trial type by constructing kernel density estimates of the observed responses. For lead and test trials, which do not lead to confusion, the density estimate was based directly on the observed distributions. For trail trials, to account for confusion, we first filtered the trials such that only those with preceding test trials in region λ were included.

**Fig S4.**
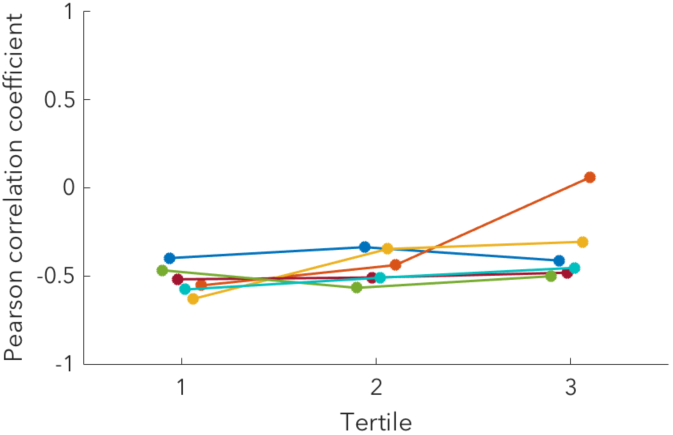
Estimates of the parameters *γ* and *σ* are moderately anticorrelated. We determined the linear correlation between estimates of the parameters *γ* and *σ* by calculating the Pearson correlation coefficient. We calculated this coefficient separately for each subject and for each tertile and typically found a negative value between −0.3 and −0.7, indicating moderate anticorrelation.

## Acknowledgments

We would like to thank Wittawat Jitkrittum, Jesse Geerts, Tian Tian and Li Wenliang for fruitful discussions. Steve Cabilio developed and maintained the experimental-control and data acquisition software used in this study. The experimental-control and data acquisition hardware was designed, built, and maintained by David Munro.

